# Rapid screening of pest resistance genes in maize using a sugarcane mosaic virus vector

**DOI:** 10.1101/2021.01.13.425472

**Authors:** Seung Ho Chung, Mahdiyeh Bigham, Ryan R. Lappe, Barry Chan, Ugrappa Nagalakshmi, Steven A. Whitham, Savithramma P. Dinesh-Kumar, Georg Jander

## Abstract

*Spodoptera frugiperda* (fall armyworm) is a notorious pest that threatens maize production world-wide. Current control measures involve the use of chemical insecticides and transgenic maize expressing *Bacillus thuringiensis* (*Bt*) toxins. Although several additional transgenes have confirmed insecticidal activity in other plants, limited research has been conducted in maize, at least partially due to the technical difficulty of maize transformation. Here, we describe implementation of a sugarcane mosaic virus (SCMV) vector for rapidly testing the efficacy of transgenes for the control of *S. frugiperda* in maize. Four categories of proteins were tested using the SCMV vector: (i) maize defense signaling proteins: peptide elicitors (Pep1 and Pep3) and jasmonate acid conjugating enzymes (JAR1a and JAR1b); (ii) maize defensive proteins: the previously identified ribosome-inactivating protein (RIP2) and maize proteinase inhibitor (MPI), and two proteins with predicted but unconfirmed anti-insect activities, an antimicrobial peptide (AMP) and a lectin (JAC1); (iii) lectins from other plant species: *Allium cepa* agglutinin (ACA) and *Galanthus nivalis* agglutinin (GNA); and (iv) spider and scorpion toxins: peptides from *Urodacus yaschenkoi* (UyCT3 and UyCT5) and *Hadronyche versuta* (Hvt). In most cases, *S. frugiperda* larval growth on maize was reduced by transient SCMV-mediated overexpression of genes encoding these proteins. Additionally, experiments with some of the SCMV-expressed genes showed effectiveness against two aphid species, *Rhopalosiphum maidis* (corn leaf aphid) and *Myzus persicae* (green peach aphid). Together, these results demonstrate that SCMV vectors can be exploited as a rapid screening method for testing the efficacy and insecticidal activity of candidate genes in maize.

## Introduction

Maize (*Zea mays*) is one of the world’s most important cereal crops, serving not only as a food source for humans and livestock, but also as a raw material for the production of ethanol and other industrial products (Ai and Jane, 2016; Chaudhary *et al*., 2014). The needs of an ever-expanding population will lead to increasing demands on maize production in the coming years. Therefore, maintaining adequate maize yields will require reducing not only reducing the cost of agricultural inputs, but also the negative impacts of biotic and abiotic stresses that that limit maize productivity.

More than 100 species of insect pests limit maize productivity in agricultural fields (McMullen *et al*., 2009; Meihls *et al*., 2012). Among these pests, one of the most damaging is *Spodoptera frugiperda* (fall armyworm; Figure 1a), a lepidopteran species that is indigenous to the Americas but recently has become invasive in Africa and Asia (Food and Agriculture Organization of the United Nations, 2018; Goergen *et al*., 2016). By consuming all above-ground plant parts, *S. frugiperda* larvae reduce photosynthetic area, cause developmental delays, and decrease yield.

**Figure 1.**
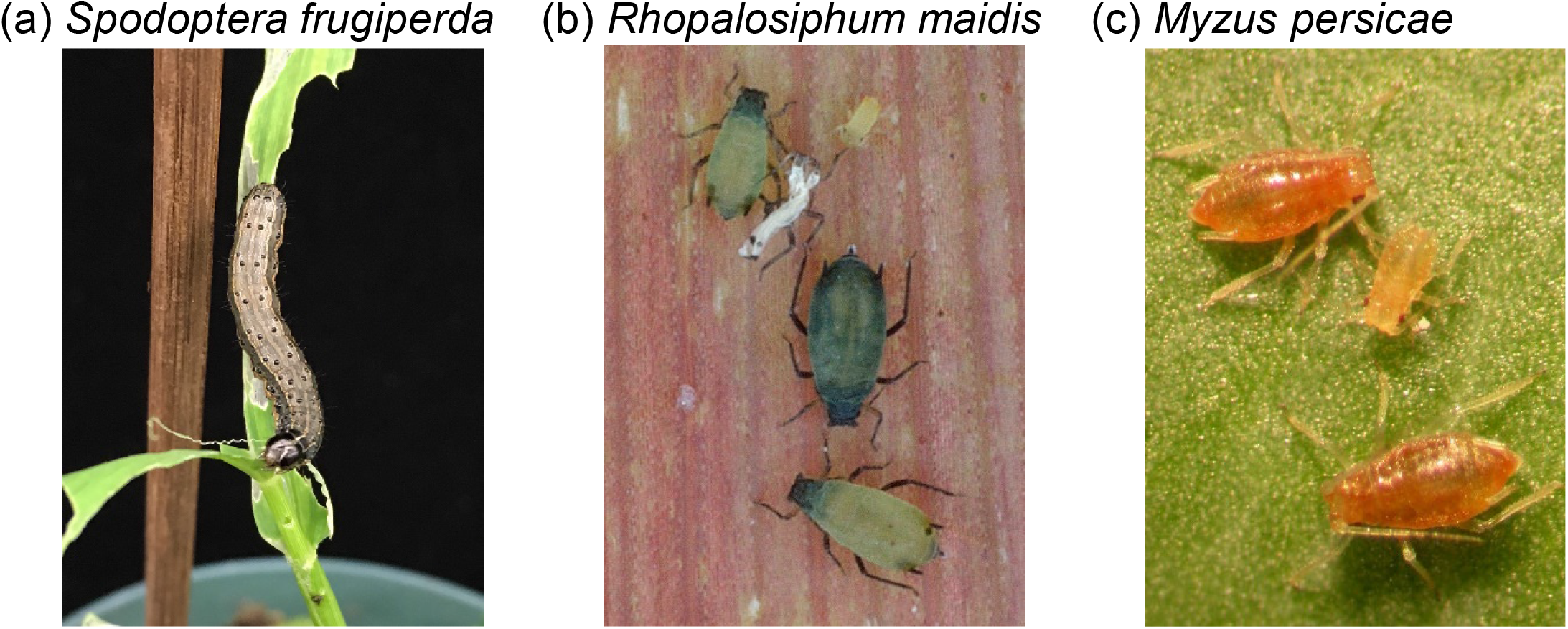
Insect species used in this study. (a) *Spodoptera frugiperda*, fall armyworm, (b) *Rhopalosiphum maidis*, corn leaf aphid, (c) *Myzus persicae*, green peach aphid.

Currently available *S. frugiperda* control methods, both application of chemical insecticides (Togola *et al*., 2018) and transgenic maize producing *Bacillus thuringiensis* (*Bt*) toxins (Huang *et al*., 2014; Tabashnik and Carrière, 2017), are becoming less effective as the insects develop resistance. Therefore, there is a need to screen for additional transgenes that can be used to enhance maize resistance to *S. frugiperda* feeding. Broadly, such approaches can include upregulation of maize defense signaling, overexpression of individual maize defensive proteins, and expression of exogenous insecticidal proteins.

Plant elicitor peptides (Peps) trigger anti-herbivore defense responses (Huffaker, 2015; Huffaker *et al*., 2013; Poretsky *et al*., 2020). In maize, *Zm*Pep1 and *Zm*Pep3 upregulate defenses, at least in part by induction of the jasmonic acid (JA) signaling pathway (Huffaker *et al*., 2013, 2011). A key step in the JA pathway is the conjugation of JA with isoleucine by JAR1 (JASMONATE RESISTANT 1) enzymes (Koo and Howe, 2009; Staswick *et al*., 2002) to form JA-isoleucine. Expression of *JAR1a* and *JAR1b*, two of the five predicted *JAR* genes in maize (Borrego and Kolomiets, 2016), is highly induced by *Spodoptera exigua* (beet armyworm) herbivory (Tzin *et al*., 2017). Thus, these maize genes are good targets for overexpression to enhance resistance against *S. frugiperda*.

Maize and other plants produce ribosome-inactivating proteins (RIPs) that block ribosome function by depurinating a specific adenine residue of the large ribosomal RNA (Bass *et al*., 2004; Zhu *et al*., 2018). These proteins, which are toxic for a variety of insects, including Lepidoptera (Shahidi-Noghabi *et al*., 2009) and Hemiptera (Hamshou *et al*., 2016), have been used previously in transgenic approaches. For instance, the expression of a maize kernel RIP1 in *Nicotiana tabacum* (tobacco) increased resistance to *Helicoverpa zea* (corn earworm) feeding (Dowd *et al*., 2003). The *RIP2* gene is expressed in all maize tissues except the kernels (Bass *et al*., 2004). *RIP2* expression was induced by *S. frugiperda* herbivory and recombinant RIP2 protein decreased caterpillar growth on artificial diet (Chuang, Herde, *et al*., 2014).

Two additional classes of maize proteins with anti-herbivore activity are proteinase inhibitors and antimicrobial peptides (Campos *et al*., 2018; Koiwa *et al*., 1997). Proteinase inhibitors, which are produced by many plant families, impair the growth and survival of insects by disrupting the function of digestive enzymes. Maize proteinase inhibitor (*MPI*) expression was induced by both caterpillar herbivory and JA signaling (Cordero *et al*., 1994; Shivaji *et al*., 2010; Tamayo *et al*., 2000). Heterologous expression of *MPI* in rice increased resistance to *Chilo suppressalis* (striped stem borer) (Vila *et al*., 2005). Cyclotides are macrocyclic insecticidal peptides with the length of about 30 amino acids and a conserved cystine knot motif containing three disulfide bonds (Campos *et al*., 2018; Craik *et al*., 1999; Weidmann and Craik, 2016). Cyclotide Kalata B1 from *Oldenlandia affinis* decreased the growth of *Helicoverpa armigera* (corn earworm) larvae by rupturing epithelial cells in the midgut (Barbeta *et al*., 2008). Cycloviolacins, cyclotides from *Viola odorata*, negatively affected the probing and feeding behavior of *Mysus persicae* (green peach aphid), suggesting that cycloviolacins limit aphid population growth (Dancewicz *et al*., 2020). Among predicted antimicrobial peptides in maize, a few belong to the cyclotide family (Mulvenna *et al*., 2006; Noonan *et al*., 2017), but their efficacy against insects has not been confirmed.

Plant lectins, carbohydrate-binding proteins that interact with glycoproteins and glycan structures in insect guts, have antinutritional or insecticidal effects (Macedo *et al*., 2015). For instance, snowdrop lectin (*Galanthus nivalis* agglutinin; GNA), onion lectin (*Allium cepa* agglutinin; ACA) and garlic (*Allium sativum*) leaf lectin reduce nutrient uptake and growth in wide range of insects (Vandenborre *et al*., 2011). Expression of a maize lectin gene, Jacalin 1 (*JAC1*), is induced by JA, an indication that it may provide protection against herbivory (Van Damme *et al*., 2004). There can be additive or even synergistic effects if lectins are co-expressed or fused to scorpion or spider venom peptides. For instance, the insecticidal efficacy of GNA was increased by fusions to Hvt (Fitches *et al*., 2012), ButaIT from *Mesobuthus tamulus* (Fitches *et al*., 2010), AaIT from *Androctonus australis* (Liu *et al*., 2016), and δ-amaurobitoxin-PIla from *Pireneitega luctuosus* (Yang *et al*., 2014).

Scorpion and spider venoms, which contain numerous insecticidal toxins (King and Hardy, 2013; Ortiz *et al*., 2015), have been explored as sources of insecticidal peptides. UyCT3 and UyCT5, two antimicrobial peptides that are produced in the venom glands of *Urodacus yaschenkoi* (inland robust scorpion) decrease the fitness of *Acyrthosiphon pisum* (pea aphid) and the density of the primary symbiont, *Buchnera aphidicola*, suggesting that those are promising candidates for the production of insect-resistant transgenic plants (Luna-Ramirez *et al*., 2017). Similarly, ω-hexatoxin Hv1a (Hvt, also called ω-ACTX Hv1a) from *Hadronyche versuta* (Blue Mountains funnel web spider) is broadly effective against both lepidopteran and hemipteran pests when expressed in transgenic plants (Javaid *et al*., 2016; Ullah *et al*., 2015; Rauf *et al*., 2019). Given the broad insecticidal activity of these spider and scorpion venom proteins, we hypothesized that they would also be effective in reducing *S. frugiperda* growth.

Testing the effectiveness of transgenes for controlling pest insects on maize is limited by the high cost of maize transformation and the often greater than one-year timeline that is required to obtain transgenic maize plants for experiments. Therefore, we are proposing an alternate approach, whereby the efficacy of transgenes that enhance maize pest tolerance is tested by transient expression in maize using a virus vector. Sugarcane mosaic virus (SCMV), a positive-sense single-stranded RNA virus, which has been adapted for efficient transgene expression in maize (Mei *et al*., 2019), is attractive vector for such experiments. Genes of interest are inserted between the SCMV *P1* and *HC-Pro* cistrons in SCMV-CS3, a newly created plasmid vector, maize seedlings are infected by particle bombardment or *Agrobacterium* inoculation, and the pest resistance of the infected plants can be assessed after three weeks. Here we show that virus-mediated expression of maize defense-regulating proteins, maize insecticidal proteins, and exogenous toxins can reduce the growth of insect pests. We demonstrate the insect-controlling properties of not only known proteins but also two maize proteins that were not previously confirmed to have anti-herbivore properties. Furthermore, we show that expression of transgenes using SCMV also is effective in reducing production by two hemipteran pests, *Rhopalosiphum maidis* (corn leaf aphid; Figure 1b) and *M. persicae* (Figure 1c).

## Materials and methods

### Plants and insects

Maize (*Zea mays*) plants, sweet corn variety Golden Bantam (West Coast Seeds, British Columbia, Canada) and inbred lines P39 and B73, were grown in a maize mix [0.16 m^3^ Metro-Mix 360 (Scotts, Marysville, OH, USA), 0.45 kg finely ground lime, 0.45 kg Peters Unimix (Griffin Greenhouse Supplies, Auburn, NY, USA), 68 kg Turface MVP (Banfield-Baker Corp., Horseheads, NY, USA), 23 kg coarse quartz sand, and 0.018 m^3^ pasteurized field soil]. All plants, including those used for SCMV propagation and insect bioassays, were maintained in a growth chamber at 23°C with a 16:8 light:dark cycle. Unless specified otherwise, Golden Bantam maize was used for the described experiments.

Eggs of *S. frugiperda* (fall armyworm) were purchased from Benzon Research (Carlisle, PA, USA) and maintained on an artificial diet (Fall Armyworm Diet, Southland Products Inc, Lake Village, AR, USA) in an incubator at 28°C. A colony of a genome-sequenced *R. maidis* lineage (W. Chen *et al*., 2019) was maintained on maize (Golden Bantam or P39) and a colony of a previously described tobacco-adapted strain of *M. persicae* (Ramsey *et al*., 2007) was maintained on *Nicotiana tabacum* (tobacco) at 23°C under 16:8 light:dark cycle. Both *R. maidis* and *M. persicae* were originally collected in USDA-ARS greenhouses by Steward Gray (Robert W. Holley Center for Agriculture & Health, Ithaca, NY, USA).

### Cloning of candidate genes into *Sugarcane mosaic virus* for protein expression

The pSCMV-CS3 expression vector used in this work was derived from pSCMV-CS2 (Mei *et al*., 2019), which was modified to contain the CS3 restriction sites in the MCS between the P1 and HC-Pro cistrons (Figure 2a). The modified pSCMV-CS2 genome plus the flanking 35S promoter and NOS terminator was amplified with SuperFi polymerase (ThermoFisher Scientific, Waltham, MA) using primers DCPacI 1380 F and DCPacI 1380 R (Supplementary Table S1). The pCAMBIA1380 backbone (www.cambia.org) was amplified with SuperFi polymerase using primers 1380F and 1380R (Supplementary Table S1). The two PCR fragments were subsequently assembled into pCAMBIA1380-SCMV-CS3, hereafter referred to as pSCMV-CS3, by Gibson Assembly (New England Biolabs, Ipswich, MA). pSCMV-GFP was created by amplifying the mEGFP coding sequence (Zacharias *et al*., 2002) using SuperFi polymerase with the GFP-Psp and GFP-SbfI primers (Supplementary Table S1). The resulting amplicon was digested with *Psp*OMI and *Sbf*I, gel purified and ligated into similarly digested pSCMV-CS3.

**Figure 2.**
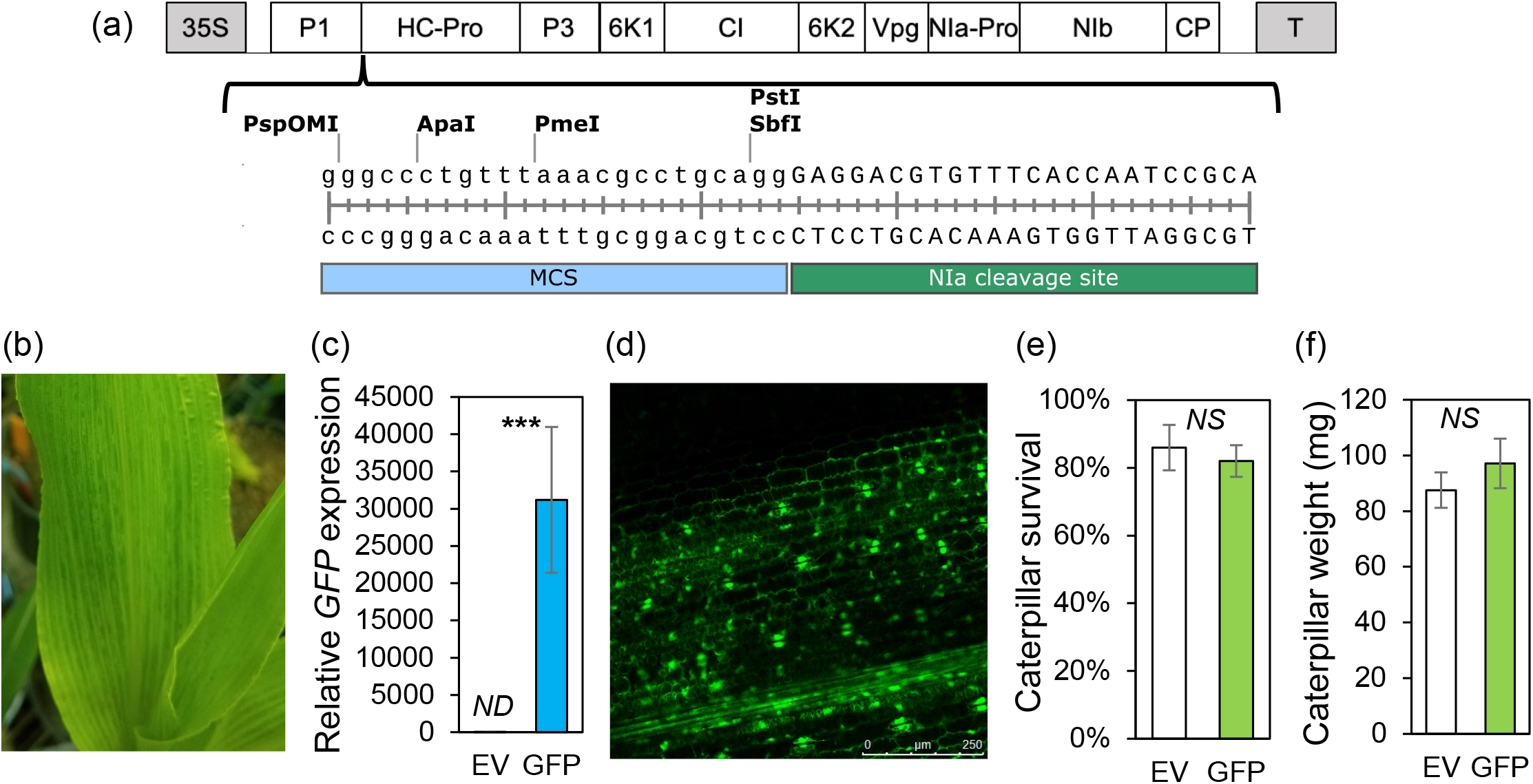
Use of a sugarcane mosaic virus (SCMV) to transiently express *GFP* in maize. (a) Schematic diagram of the SCMV-CS3 cloning vector. A multiple cloning site (MCS) was inserted between P1 and HC-Pro genes of SCMV. 35S: cauli?ower mosaic virus 35S promoter; T: NOS terminator. (b) Mosaic infection symptom of SCMV vector in maize three weeks post inoculation. (c) Transient overexpression of *GFP* in plants infected by SCMV-empty vector (EV) or -*GFP*. Gene expression was determined by qRT-PCR three weeks post-inoculation. Means +/− s.e. of N = 5, ***P < 0.001, *t-*test. (d) Confocal image of foliar GFP expression. (e, f) *S. frugiperda* survival and larval growth on plants infected by EV or SCMV-*GFP*. Two-day-old caterpillars were fed on infected plants for one week and caterpillar mass was determined. Means + s.e. of N = 10, *NS*= non-significant, *ND* = not detected, EV = empty vector, MCS = multiple cloning site.

Genes encoding maize defense regulators and protein toxins were amplified from the B73 cDNA template with gene-specific primers containing restriction sites at the 5’ end for cloning into SCMV vector. PCR products were gel purified, digested with the corresponding restriction enzymes and cloned into the SCMV vector (Figure 2a). The *UyCT3* and *UyCT5* genes (codon optimized for maize), as well as the maize *Pep1* and *Pep3* genes, were synthesized by Genscript Biotech (Piscataway, NJ, USA) and cloned into the SCMV vector. To generate SCMV with *Hvt, ACA, GNA, RIP2, Hvt-ACA, Hvt*\*-*ACA, RIP2-GNA*, and *Hvt-GNA*, primers listed in Supplementary Table S1 and DNA synthesized by Genewiz (South Plainfield, NJ, USA) were used as template and the resulting PCR products were gel purified and cloned into PspOMI-SbfI cut SCMV by following the NEBuilder HiFi DNA Assembly method (NEB, Ipswich, MA, USA). Primers, restriction sites, and cloning methods used in this study are listed in Supplementary Table S1.

### Inoculation of maize with SCMV constructs

SCMV constructs were delivered into maize plants by particle bombardment using a Biolistic PDS-1000/He system (Bio-Rad, Hercules, CA, USA) as described previously (Mei and Whitham, 2018). One µg of the plasmid DNA was coated on 3 mg 1.0 µm diameter gold particles, and the coated gold particles were distributed onto five microcarriers and allowed to air dry. Plants were placed in the dark 12 h before the particle bombardment. Two leaves of one-week-old plants were bombarded with using 1,100 psi rupture disks at a distance of 6 cm (between stopping screen and leaves).

Agroinoculation was used to initiate maize infections with the SCMV constructs containing *Hvt, ACA*, and *UyCT3*. The constructs were transformed into the *Agrobacterium tumefaciens* strain GV3101 and an *A. tumefaciens* suspension with optical density at 600 nm (OD_600_) = 1.0 in infiltration buffer (200 μM acetosyringone, 10 mM MES, pH 5.6, and 10 mM MgCl_2_) was injected above the coleoptile node of one-week-old plants.

After the initial infection by particle bombardment or Agroinoculation, SCMV constructs were further propagated by rub-inoculation. Leaf sap of SCMV-infected plants was prepared by grinding 0.5 g leaf tissue in 5 ml of 50 mM pH 7.0 potassium phosphate buffer. One-week-old maize plants were dusted with 600-mesh carborundum and mechanically inoculated by rubbing leaf sap from virus-infected maize plants on two leaves.

### Confocal microscopy

Three weeks post-inoculation, leaf samples were collected from the seventh or eighth leaves of maize plants infected with SCMV-*GFP*. The samples were observed at an excitation of 488 nm. The emitted fluorescence signal was monitored from 505 to 545 nm using a SP5 Leica Confocal Microscope in the Plant Cell Imaging Center of Boyce Thompson Institute. A scan of fluorescence across a range of wavelengths (lambda scan) was used to confirm that the observed signal was derived from GFP rather than endogenous maize fluorescence.

### Insect bioassays

To determine the effect of defensive proteins on the growth of *S. frugiperda*, SCMV-infected plants, three weeks post-inoculation, were used for caterpillar bioassays. Five two-day-old caterpillars were placed on each plant and enclosed using perforated plastic bags (13 cm x 61 cm, https://www.clearbags.com). The caterpillar fresh mass was measured one week later. For aphid bioassays, eight 10-day-old apterous adult *R. maidis* or ten 10-day-old apterous adult *M. persicae* were placed on each virus-infected plant 15-18 days after SCMV inoculation and enclosed using perforated plastic bags. The total numbers of aphids were counted one week later.

### RNA extraction, cDNA synthesis, RT-PCR, and quantitative real-time PCR (qRT-PCR)

Three weeks post-inoculation and prior to the insect bioassays, leaf tissue was collected from the seventh or eighth leaves of infected plants, flash-frozen in liquid nitrogen, and stored at −80 °C. RNA was extracted using TRIzoL Reagent (Invitrogen, Carlsbad, CA, USA) and treated with RQ1 RNase-free DNase (Promega, Madison, WI, USA). One microgram of RNA was used to synthesize first-strand cDNA using High-Capacity cDNA Reverse Transcription Kit (Applied Biosystems, Foster City, CA, USA) with random primers. To verify the expression of insect resistance genes cloned into the SCMV vector, qRT-PCR was conducted with gene-specific primers (Supplementary Table S1). For the inserts of less than 75 bp (*Pep1, Pep3, UyCT3*, and *UyCT5*), one primer was designed to bind to the insert and another primer was designed to bind to the region flanking the cloning site, with the amplification products ranging in size from 100-150 bp. The reactions consisted of 5.0 µl of the PowerUp SYBR Green PCR master mix (Applied Biosystems), 0.6 µl primer mix (300 nM for the final concentration of each primer) and 2 µl of cDNA (1:10 dilution with nuclease-free H_2_O) in 10 µl total volume. Template-free reactions were included as negative controls. The PCR amplification was performed on QuantStudio 6 Flex Real-Time PCR Systems (Applied Biosystems, Foster City, CA, USA) with the following conditions: 2 min at 50°C, 2 min at 95°C, 40 cycles of 95°C for 15 sec and 60°C for 1 min. Primer specificity was confirmed by melting curve analysis. Mean cycle threshold values of duplicates of each sample were normalized using two reference genes, *Actin* and *EF1-α*. Relative gene expression values were calculated using 2^-ΔΔCt^ method (Livak and Schmittgen, 2001).

### Statistical analysis

All statistical analyses were conducted using R (R Core Team, 2017). Data for gene expression and larval mass of *S. frugiperda* larvae, and aphid fecundity were compared using analysis of variance (ANOVA) followed by Tukey’s test or Dunnett’s test relative to the GFP control or *t*-test. Gene expression data were log2 transformed before the statistical analysis to meet the assumptions of ANOVA but untransformed data are presented in the figures. Survival of *S. frugiperda* larvae was analyzed using non-parametric Kruskal-Wallis tests. Raw data underlying the bar graphs are presented in Supplemental Tables S2-S8.

## Results

### SCMV-GFP does not affect *S. frugiperda* growth on maize

A previously described SCMV cloning vector (Mei *et al*., 2019) was modified to produce SCMV-CS3 (Figure 2a) and the mEGFP coding sequence (Zacharias *et al*., 2002) was placed in the multiple cloning site to produce SCMV-*GFP*. An empty vector control virus (SCMV-EV) and SCMV-*GFP* were used to inoculate one-week-old maize seedlings. Both SCMV-*GFP* and SCMV-EV caused mosaic symptoms within two weeks after inoculation and continued to spread in newly emerging leaves of the infected plants (Figure 2b). Three weeks post-inoculation, the *GFP* expression level was significantly higher in plants infected by SCMV-*GFP* than SCMV-EV (Figure 2c). Infected leaves were examined using confocal microscopy and the green fluorescence signal was only detected in leaves infected with SCMV-*GFP* (Figure 2d). Neither *S. frugiperda* larval survival (Figure 2e) nor larval growth (Figure 2f) differed significantly between plants infected by SCMV-*GFP* and SCMV-EV. SCMV-*GFP* was used as a transgene-expressing virus control treatment in subsequent experiments to test the efficacy of insect growth-inhibiting proteins (Table 1) against *S. frugiperda*, *R. maidis*, and *M. persicae* (Figure 1a-c).

**Table 1.** Insect resistance genes that were tested in maize by transience expression using *Sugarcane mosaic virus*

| Source species | Gene name | GenBank ID | MaizeGDB ID | Description |
| --- | --- | --- | --- | --- |
| *Zea mays* | *Pep1* | XM_008670579 | GRMZM2G055447 | Peptide elicitor 1 |
| *Zea mays* | *Pep3* | XM_008670581 | GRMZM2G339117 | Peptide elicitor 3 |
| *Zea mays* | *JAR1a* | NM_001174342 | GRMZM2G091276 | Jasmonate-isoleucine conjugating enzyme |
| *Zea mays* | *JAR1b* | NM_001361064 | GRMZM2G162413 | Jasmonate-isoleucine conjugating enzyme |
| *Zea mays* | *RIP2* | NM_001137489 | GRMZM2G119705 | Ribosome-inactivating protein 2 |
| *Zea mays* | *JAC1* | NM_001148875 | GRMZM2G050412 | Jacalin 1, maize lectin |
| *Zea mays* | *AMP* | NM_001371023 | GRMZM2G032198 | Cyclotide antimicrobial peptide |
| *Zea mays* | *MPI* | X78988 | GRMZM2G028393 | Maize proteinase inhibitor |
| *Allium cepa* | *ACA* | DQ255944 |  | Onion lectin |
| *Galanthus nivalis* | *GNA* | M55556 |  | Snowdrop lectin |
| *Hadronyche versuta* | *Hvt* | AJ938032 | | ω-hexatoxin-Hv1a, spider venom toxin |
| *Urodacus yaschenkoi* | *UyCT3* | JX274241 |  | Scorpion venom toxin, antimicrobial peptide |
| *Urodacus yaschenkoi* | *UyCT5* | JX274242 |  | Scorpion venom toxin, antimicrobial peptide |

### Expression of endogenous maize genes using SCMV enhances *S. frugiperda* resistance

We cloned the 69 bp sequences of the maize *Pep1* and *Pep3* defense elicitors (Huffaker *et al*., 2011, 2013) into pSCMV-CS3 and inoculated seedlings of sweet corn variety Golden Bantam. *Pep1* and *Pep3* expression was confirmed by qRT-PCR three weeks post-inoculation (Figure 3a,b). Relative to SCMV-*GFP*, infection with SCMV-*Pep1* and SCMV-*Pep3* caused increased transcript accumulation of maize proteinase inhibitor (*MPI*; Figure 3c), a JA pathway marker gene with antiherbivore activity (Chuang, Ray, *et al*., 2014). *Pep1* and *Pep3* expression decreased the growth of *S. frugiperda* larvae on Golden Bantam maize by 25% and 51%, respectively, compared to GFP control plants (Figure 3d), but did not affect survival of the larvae (Table S4). Similar results were obtained for *Pep3* using popcorn (P39) and field corn (B73) inbred lines (Figure 3e,f), showing that the effect on *S. frugiperda* growth is not specific to Golden Bantam.

**Figure 3.**
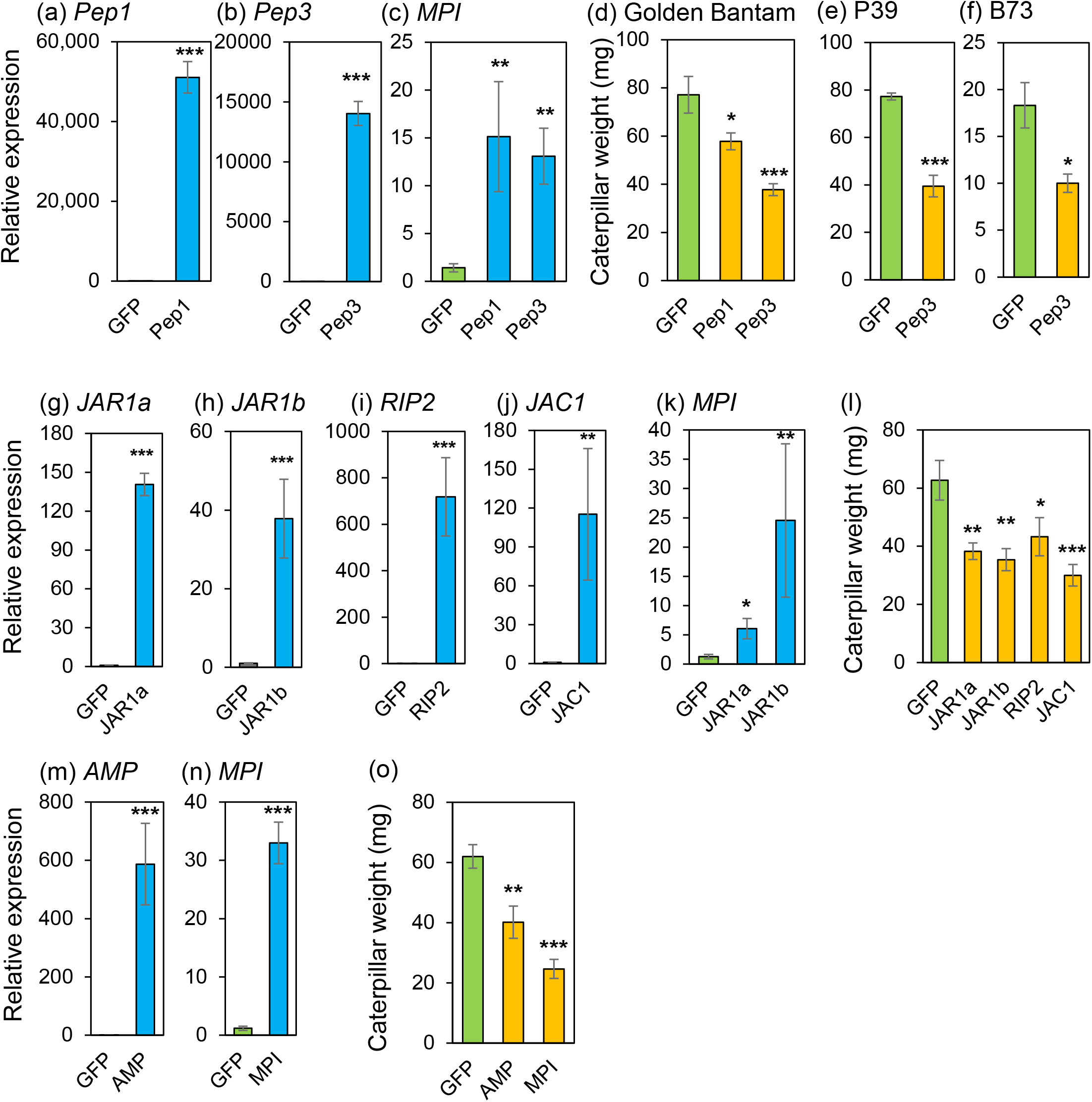
Gene expression and *Spodoptera frugiperda* larval growth on plants expressing maize defense genes. (a,b) Transient overexpression of *Pep1* and *Pep3* in Golden Bantam plants infected by SCMV-*GFP*, -*Pep1* or - *Pep3*. (c) The expression level of *MPI* in plants expressing SCMV-encoded *GFP, Pep1*, or *Pep3*. Gene expression was determined by qRT-PCR three weeks post inoculation. (d) Performance of *S. frugiperda* on Golden Bantam plants expressing *GFP, Pep1*, or *Pep3*. (e,f) Performance of *S. frugiperda* on P39 and B73 plants expressing *GFP* or *Pep3*. (g-j, m,n) Transgene expression in Golden Bantam plants infected with SCMV-*GFP*, -*JAR1a*, -*JAR1b*, -*RIP2*, -*JAC1, AMP, or MPI*. (k) The expression level of *MPI* in Golden Bantam plants expressing *GFP, JAR1a*, or *JAR1b*. (l, o) The performance of *S. frugiperda* on plants expressing *GFP, JAR1a, JAR1b, RIP2, JAC1, AMP, or MPI*. Caterpillars were confined on infected plants three weeks post inoculation and mass was measured one week later. Means +/− s.e. of N = 5 for gene expression; N = 10-12 for insect bioassays, *P < 0.05; **P < 0.01; ***P < 0.001 relative to GFP control, Dunnett’s test.

We targeted previously investigated maize insect resistance genes (*JAR1a, JAR1b, RIP2*, and *MPI*) and two predicted resistance genes (*AMP* and *JAC1*) for overexpression using SCMV. A previously uncharacterized gene (GRMZM2G032198) encoding a maize cyclotide antimicrobial peptides was designated as *AMP*. Expression of both *AMP* and *JAC1* was induced by *S. frugiperda* herbivory (Figure S1), suggesting that these genes are involved in maize insect resistance. The expression levels of *JAR1a, JAR1b, RIP2, JAC1, AMP*, and *MPI* were significantly higher in plants infected by the corresponding SCMV constructs than in SCMV-*GFP* control plants (Figure 3g-j,m,n). SCMV-*JAR1a* and SCMV-*JAR1b* also significantly increased *MPI* expression (Figure 3k), confirming upregulation of JA-related defense pathways. More importantly, the growth of *S. frugiperda* larvae was up to 60% lower on plants expressing *JAR1a, JAR1b, RIP2, JAC1, AMP*, and *MPI* than on Golden Bantam control plants infected with SCMV-*GFP* (Figure 3l,o).

### Expression of scorpion insecticidal proteins reduces *S. frugiperda* growth

To determine whether SCMV can be used to express heterologous insect resistance genes in maize, we cloned *UyCT3* and *UyCT5*, which encode *U. yaschenkoi* venom toxins, into the SCMV vector. Transgene expression in maize was confirmed by qRT-PCR (Figure 4a,b). Both SCMV-*UyCT3* and SCMV-*UyCT5* reduced *S. frugiperda* weight on P39 and B73 plants compared to SCMV-*GFP* control plants (Figure 4c,d).

**Figure 4.**
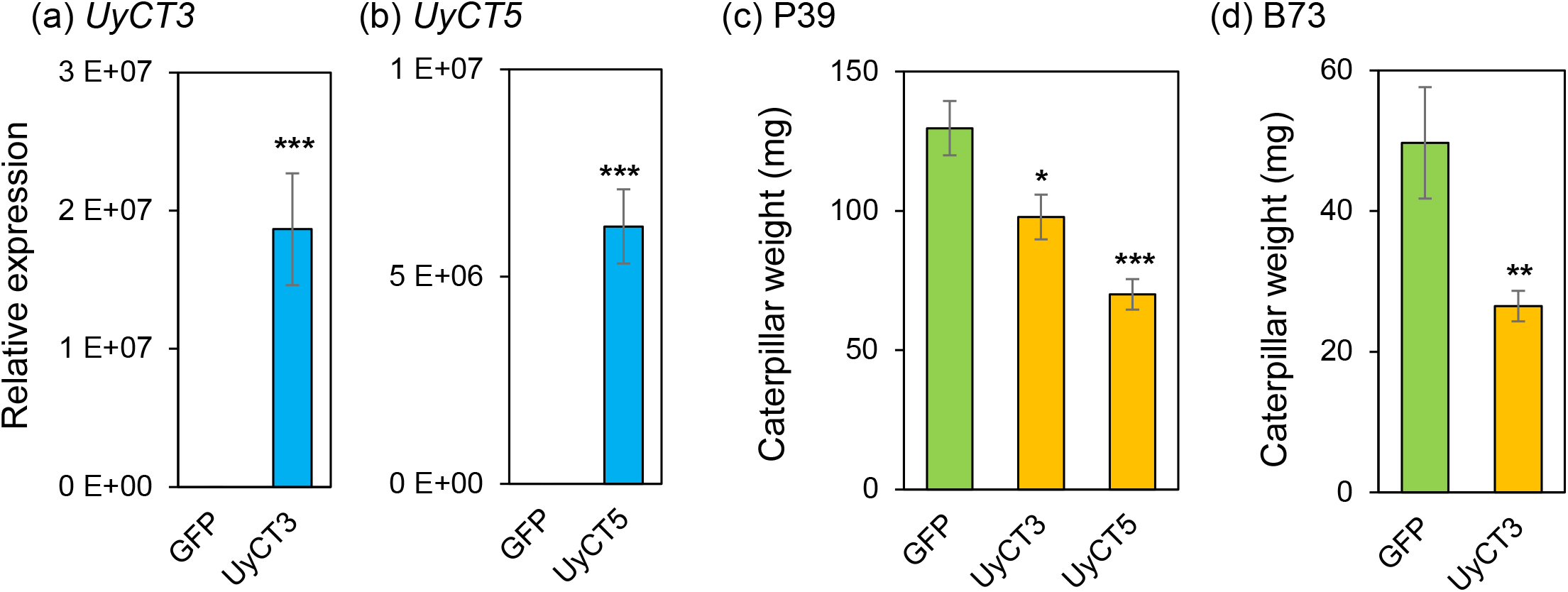
Gene expression and larval growth on P39 plants expressing heterologous insecticidal genes. (a, b) Transient overexpression of *UyCT3* and *UyCT5* in P39 plants infected by SCMV-*GFP*, -*UyCT3 or* - *UyCT5*. Gene expression was determined by qRT-PCR there weeks post inoculation. (c) The performance of *S. frugiperda* on P39 plants expressing *GFP, UyCT3* or *UyCT5*. (d) The performance of *S. frugiperda* on B73 plants expressing *GFP* or *UyCT3*. Five caterpillars were confined on infected plants three weeks post inoculation. Caterpillar weight was measured one week later. Means +/− s.e. of N = 5 for gene expression; N = 7-14 for insect bioassays, *P < 0.05; **P < 0.01; ***P < 0.001 relative to GFP control, Dunnett’s test.

### Expression of fusion proteins using SCMV has additive effects on *S. frugiperda*

Fusion of spider and scorpion neurotoxins with plant lectins can improve their toxicity (Fitches *et al*., 2012; Liu *et al*., 2016; Rauf *et al*., 2019). We investigated this effect in maize using SCMV constructs. Because SCMV vectors produce a polyprotein precursor from which functional proteins are cleaved by the NIa protease (Mei *et al*., 2019), we generated SCMV constructs with and without an NIa cleave site between the venom toxin and lectin to determine whether the proteins are more efficacious separately or as fusion proteins. The expression levels of *H. versuta* toxin (*Hvt*) and *A. cepa* agglutinin (*ACA*) genes were significantly higher in plants infected by each corresponding SCMV constructs than SCMV-*GFP* control (Figure 5a and 5b), and the presence of protease cleavage site between *Hvt* and *ACA* did not affect *Hvt* and *ACA* transcript accumulation. Expression of single proteins did not significantly decrease *S. frugiperda* growth relative to the GFP control (Figure 5c). However, the expression of fusion protein by SCMV-*Hvt*-*ACA* or two individual proteins of Hvt and ACA by SCMV-*Hvt*-cleavage site-*ACA* reduced the larval growth by 39% and 46%, relative to the GFP control (Figure 5c), respectively, suggesting an additive effect from the expression the two genes.

**Figure 5.**
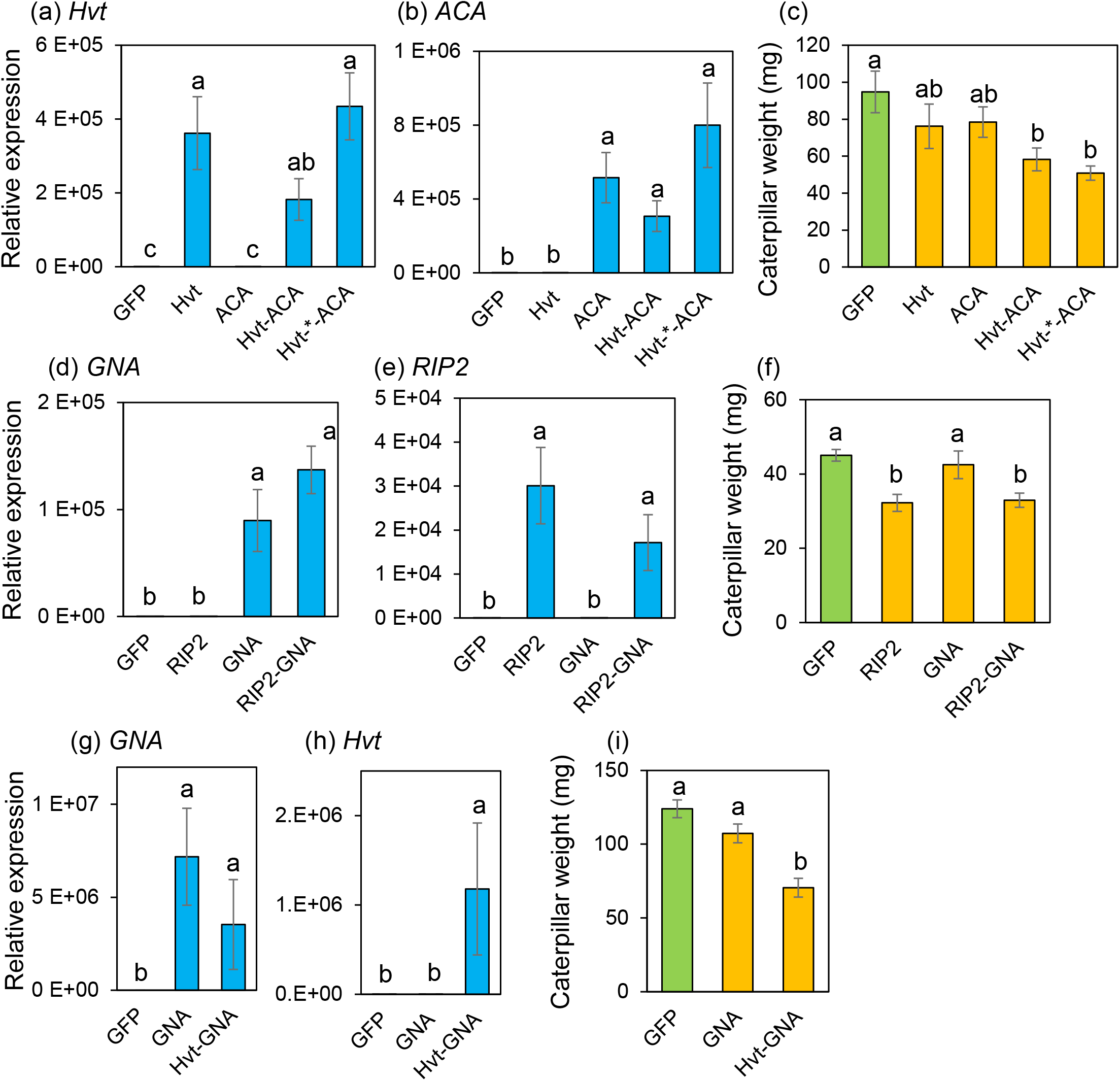
Gene expression and larval growth on plants expressing heterologous insecticidal genes. (a, b) Transient overexpression of *Hvt* and *ACA* in plants infected by SCMV-*GFP*, *-Hvt, -ACA, -Hvt-ACA* or *-Hvt-*-ACA*. * represents a protease cleavage site. (c) The performance of *Spodoptera frugiperda* on plants expressing *GFP, Hvt, ACA, Hvt-ACA* or *Hvt-*-ACA*. (d, e) Transient overexpression of *RIP2* and *GNA* in plants infected by SCMV-*GFP*, -*RIP2*, -*GNA* or -*RIP2-GNA*. (f) Performance of *S. frugiperda* on plants expressing *GFP, RIP2, GNA*, or *RIP2-GNA*. (g, h) Transient overexpression of *GNA* and *Hvt* in plants infected by SCMV-*GFP, -Hvt*, -*GNA*, or -*Hvt-GNA*. (i) Performance of *S. frugiperda* on plants expressing *GFP, Hvt, GNA*, or *Hvt-GNA*. Gene expression was determined by qRT-PCR there weeks post-inoculation. Caterpillars were confined on infected plants three weeks post-inoculation and caterpillar weight was measured one week later. Means +/− s.e. of N = 5 for gene expression and N = 10-12 for insect bioassays. Different letters indicate significant differences, P < 0.05, ANOVA followed by Tukey’s HSD test.

As the presence of a cleavage site between the two protein components did not increase efficacy (Figure 5c), we tested insecticidal activity of fusion proteins without viral protease cleavage sites in subsequent experiments. SCMV constructs were made with a maize defense gene, *RIP2*, and a spider insecticidal protein, *Hvt*, fused to *G. nivalis* agglutinin (*GNA*). We confirmed that each gene was expressed in plants infected by the corresponding construct (Figure 5d,e,g,h). Although the expression of *GNA* alone did not affect *S. frugiperda* larval growth, infection with SCMV-*RIP2* and SCMV-*RIP2-GNA* reduced *S. frugiperda* growth by 28% and 27%, respectively (Figure 5f). In the case of *Hvt* and *GNA*, neither gene by itself significantly reduced caterpillar growth. However, the SCMV-*Hvt-GNA* fusion construct significantly decreased *S. frugiperda* larval growth compared to GFP control protein (Figure 5i). Together, these results indicate that fusion proteins combining lectins and maize defense proteins or venom toxins can improve resistance against *S. frugiperda*.

### Expression of maize and scorpion genes enhances resistance to phloem-feeding herbivores

To investigate whether our SCMV constructs also provide protection against phloem-feeding insects, we conducted aphid bioassays using P39 plants infected with a subset of the previously tested constructs: SCMV-*GFP*, SCMV-*Pep3*, SCMV-*RIP2*, SCMV-*UyCT3*, SCMV-*UyCT5, SCMV-AMP*, and SCMV-*JAC1*. GFP expression did not affect aphid numbers compared to the empty vector control (Figure 6a,b). By contrast, expression of maize defense proteins and scorpion toxins significantly decreased progeny production by both *R. maidis* (Figure 6a,c) and *M. persicae* (Figure 6b,d) compared to SCMV-*GFP* control.

**Figure 6.**
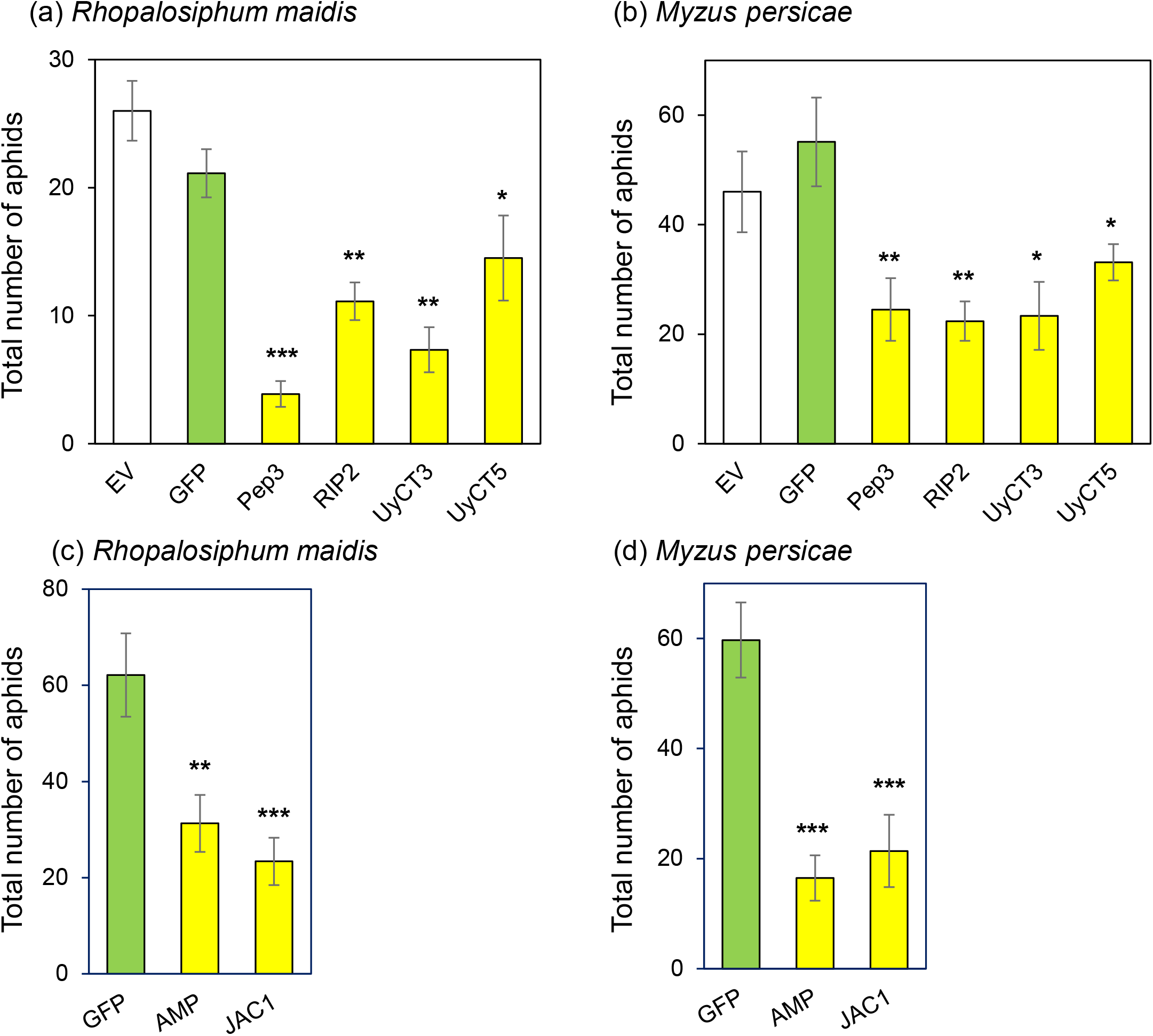
Total number of aphids on P39 plants expressing maize defense genes and scorpion toxin genes. (a,c) *Rhopalosiphum maidis* (b,d) *Myzus persicae*. Eight *R. maidis* adults or ten *M. persicae* adults were confined on plants infected by SCMV-EV, -*GFP*, -*Pep3, -RIP2, -UyCT3, UyCT5, AMP or JAC1* three weeks post inoculation. Surviving aphids and their progeny were counted one week later. Means +/− s.e. of N = 8-10, *P < 0.05; **P < 0.01; ***P < 0.001 relative to GFP control, Dunnett’s test.

## Discussion

Artificial diet assays are commonly employed for testing the oral efficacy of novel insecticidal proteins against insect herbivores (Panwar *et al*., 2018; Yao *et al*., 2003; Fitches *et al*., 2012). However, growth inhibition on artificial diet does not always correlate well with the effects that are observed when the same insecticidal proteins are subsequently expressed in transgenic plants (Khan *et al*., 2020). Both the context of the surrounding plant tissue and the localization of the insecticidal proteins in the plants could affect their toxicity against insect herbivores. Therefore, rather than pre-screening insecticidal proteins by cloning in microbial systems, purification, and artificial diet assays, we propose that transient expression using a viral vector such as SCMV will be a more effective approach for rapidly testing the *in planta* efficacy of novel insecticidal proteins.

Reduced weight gain of *S. frugiperda* in response to SCMV-mediated expression of *Pep1, Pep3, JAR1a, JAR1b, RIP2, MPI, UyCT3*, and *UyCT5* as single-gene constructs is consistent with previous reports of these genes providing protection against insect herbivory. Additionally, *AMP* and *JAC1*, two maize genes that are upregulated in response to *S. frugiperda* feeding (Figure S1), reduced caterpillar weight gain when overexpressed in maize. Although transcriptomic studies have identified numerous maize genes that are upregulated in response to arthropod feeding (Bui *et al*., 2018; Pan *et al*., 2020; Tzin *et al*., 2015, 2017; Yang *et al*., 2020; Guo *et al*., 2019; Song *et al*., 2017; Wang *et al*., 2017; Zhang *et al*., 2016), the majority of these genes have not been investigated for their role in plant defense against herbivory. This is at least in part due to the time and cost of creating maize lines that have individual genes overexpressed. Transient expression using SCMV, as we have done for *AMP* and *JAC1*, will accelerate the process of testing the defensing functions of maize genes that are induced in response to herbivory.

Although these genes increased resistance in other plant-insect studies (Liu *et al*., 2016; Ullah *et al*., 2015; Vandenborre *et al*., 2011), overexpression of *Hvt, ACA*, and *GNA* as single-gene SCMV constructs in maize did not significantly reduce *S. frugiperda* weight gain. Nevertheless, expression of gene fusions, *Hvt-ACA* and *Hvt-GNA*, reduced *S. frugiperda* weight gain, indicating that there are additive effects of the spider venom and the lectin (Figure 5c,i). This is consistent with other studies that the toxicity of spider and scorpion toxins was improved by combining with a lectin, which may facilitate the transfer of the venom proteins across the gut lumen (Fitches *et al*., 2002; Javaid *et al*., 2016; Nakasu *et al*., 2014; Rauf *et al*., 2019). In addition to these synergistic effects on larval growth, it is likely that the stacking of multiple toxic proteins with different modes of action in one viral construct will delay the development of resistance in insects (Head *et al*., 2017; Ni *et al*., 2017). In contrast to Hvt-lectin fusions, we did not observe an additive effect when RIP2 was linked to GNA (Figure 5f). This difference may be attributed to the differing origins of the toxin proteins expressed in the SCMV constructs. Whereas spider venoms like Hvt are injected directly into the hemolymph, the maize RIP2 protein would be consumed orally by lepidopteran larvae. As an endogenous maize insecticidal protein, RIP2 may bind to an as yet unknown receptor and thereby enter the midgut cells and/or the insect hemolymph. Thus, these results suggest that the fusion proteins of non-maize toxins and lectins enhance the insecticidal activity of the fusion protein.

Our observation of increased defense gene expression (Figure 3c) and reduced *S. frugiperda* weight gain (Figure 3d-f) on plants infected with SCMV-*Pep1* and SCMV-*Pep3*, are consistent with experiments showing that pre-treatment of maize plants with Pep1 and Pep3 increases JA levels, defense gene expression, and defensive metabolites, leading to reduced growth of *S. exigua* larvae (Huffaker *et al*., 2011, 2013). Under normal circumstances, peptide signaling is initiated when plasma membranes are disrupted and elicitor peptides are released from the cytoplasm into the apoplastic space (Bartels and Boller, 2015). Receptors on neighboring intact cells recognize these peptides and elicit downstream defense pathways (Lori *et al*., 2015). However, in our experiments, the expression of *MPI* was induced in plants infected by SCMV-*Pep1* and SCMV-*Pep3*, even before initiation of caterpillar feeding (Figure 3c). Pathogen attack can cause plant proteins without secretory signals to be released into apoplast (Agrawal *et al*., 2010), and is possible that SCMV infection initiated Pep1 and Pep 3 signaling in this manner

Induced defenses, in particular those regulated by the jasmonic acid pathway, typically are turned on after perception of insect herbivory by maize and other plants (Erb and Reymond, 2019; Howe and Jander, 2008). However, initiation of jasmonate-regulated defenses takes time and some lepidopteran herbivores may have the ability to suppress jasmonate signaling (C.-Y. Chen *et al*., 2019). Therefore, targeted initiation of maize defense responses by expression of regulatory proteins in SCMV may be an approach for increasing pest resistance. Such virus-mediated induction could be deployed in maize fields when there is the specific threat of insect pests such as *S. frugiperda*.

Currently available insect-resistant transgenic maize varieties, in particular those expressing *Cry, Cyt*, or *Vip* genes from *B. thuringiensis* (Bravo *et al*., 2011; Chakroun *et al*., 2016), are not effective against phloem-feeding insects such as *Rhopalosiphum maidis* (corn leaf aphid; Figure 1b) or *Myzus persicae* (green peach aphid; Figure 1c). Although the toxicity of *Bt* toxins can be increased by incorporating peptide sequences that bind to aphid guts (Chougule *et al*., 2013), this has not yet resulted in a commercially viable *Bt* toxin directed at hemipteran herbivores. Thus, there is an interest in identifying additional insecticidal proteins that can be expressed in the plant phloem to enhance aphid resistance. Our demonstration that transient expression of both endogenous maize proteins and insecticidal proteins from other species can reduce reproduction of *R. maidis* and *M. persicae* (Figure 6), suggests that SCMV-mediated overexpression can be used to rapidly screen proteins for their effectiveness against hemipteran pests of maize.

Our results demonstrate the utility of SCMV-mediated overexpression for screening the efficacy of proteins that reduce insect growth on maize plants. Virus-mediated transient expression assays included genes encoding maize regulatory proteins, endogenous maize defensive proteins, and non-maize insecticidal proteins. The main advantage of the SCMV overexpression system is a timeline that makes it possible to test the effectiveness of single- and multi-gene constructs in actual maize plants in only two months. Although the main focus of our efforts was a lepidopteran herbivore, *S. frugiperda*, we also showed efficacy of SCMV constructs against two aphid species, *R. maidis* and *M. persicae*, suggesting that the SCMV-mediated transient expression approach will be broadly useful for experiments with both chewing and piercing/sucking herbivores of maize.

## Acknowledgements

We thank Mamta Srivastava for help with confocal microscopy. This work was supported by agreement HR0011-17-2-0053 from the Defense Advanced Research Projects Agency (DARPA) Insect Allies Program with the Boyce Thompson Institute. The views and conclusions contained in this document are those of the authors and should not be interpreted as representing the official policies, either expressed or implied, of DARPA or the US Government. The US Government is authorized to reproduce and distribute reprints for Government purposes notwithstanding any copyright notation hereon.

## Short legends for Supporting Material

**Figure S1.**
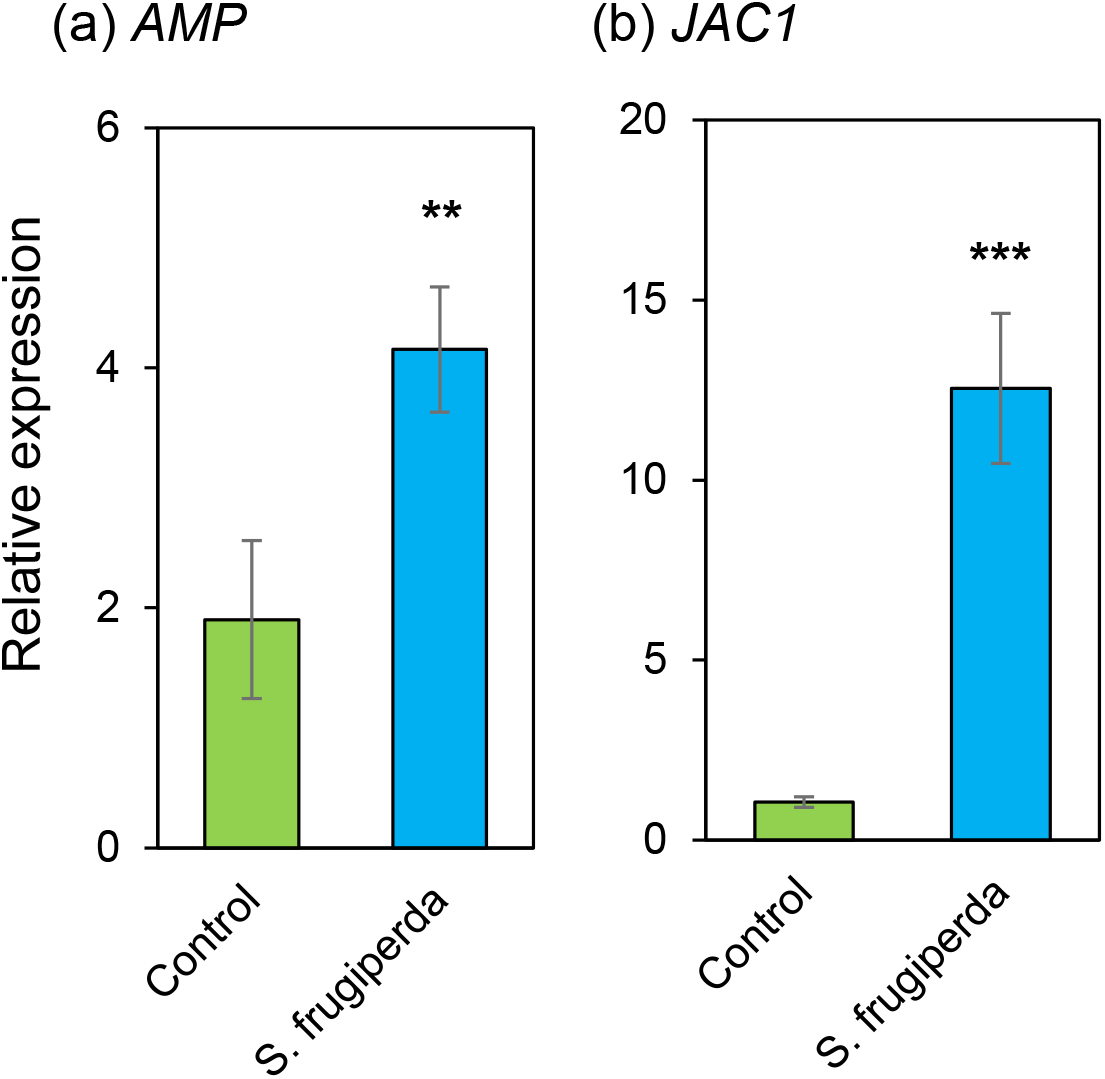
Gene expression of (a) *AMP* and (b) *JAC1* in P39 maize plants damaged by *Spodoptera frugiperda* caterpillars. Two caterpillars were confined on plants. Gene expression was determined by qRT-PCR one day after placing the caterpillars. Means +/− s.e. of N = 6, **P < 0.01; ***P < 0.001 relative to GFP control, *t-*test.

**Supplementary Table S1**. Primers used in this study.

**Supplementary Table S2**. Raw data for the graphs in Figure 2.

**Supplementary Table S3**. Raw data for the graphs in Figure 3.

**Supplementary Table S4**. Raw data for the graphs in Figure 4.

**Supplementary Table S5**. Raw data for the graphs in Figure 5.

**Supplementary Table S6**. Raw data for the graphs in Figure 6.

**Supplementary Table S7**. Raw data for the graphs in Figure S1.

